# PyLiger: Scalable single-cell multi-omic data integration in Python

**DOI:** 10.1101/2021.12.24.474131

**Authors:** Lu Lu, Joshua D. Welch

## Abstract

**Motivation:** LIGER is a widely-used R package for single-cell multi-omic data integration. However, many users prefer to analyze their single-cell datasets in Python, which offers an attractive syntax and highly-optimized scientific computing libraries for increased efficiency.

**Results:** We developed PyLiger, a Python package for integrating single-cell multi-omic datasets. PyLiger offers faster performance than the previous R implementation (2-5× speedup), interoperability with AnnData format, flexible on-disk or in-memory analysis capability, and new functionality for gene ontology enrichment analysis. The on-disk capability enables analysis of arbitrarily large single-cell datasets using fixed memory.

**Availability:** PyLiger is available on Github at https://github.com/welch-lab/pyliger and on the Python Package Index.

**Supplementary information:** Supplementary data are available at *Bioinformatics* online.

## 1 Introduction

High-throughput sequencing technologies now enable the measurement of gene expression, DNA methylation, histone modification, and chromatin accessibility at the single-cell level. Integration of such single-cell multi-omic datasets is crucial for identifying cell types and cell states across a range of biological settings. Previously, we developed LIGER (Linked Inference of Genomic Experimental Relationships), an R package that employs integrative non-negative matrix factorization to identify shared and dataset-specific factors of cellular variation (Welch *et al*., 2019). These factors then provide a principled and quantitative definition of cellular identity and how it varies across biological settings.

Many users prefer to analyze their single-cell datasets in Python, which offers an attractive syntax and highly-optimized scientific computing libraries for increased efficiency. However, there is a lack of single-cell multi-omic integration tools available in Python. The Seurat v3 (Stuart *et al*., 2019) anchors algorithm is implemented in R, as is Harmony (Korsunsky *et al*., 2019). Scanpy (Wolf *et al*., 2018) offers excellent libraries for single-cell RNA-seq analysis, including batch correction with the BBKNN algorithm, but this approach is not designed for multi-omic integration such as combining scRNA and snATAC from different cells. The scvi-tools (Gayoso *et al*., 2021) library similarly provides options for scRNA integration, but is not designed for integrating different single-cell modalities from different individual cells.

To address these limitations, we developed PyLiger, a Python implementation of LIGER.

## 2 Results

### 2.1 Python implementation of LIGER

We translated the complete, established LIGER framework into Python. Key functions includes integration of multiple single-cell datasets using integrative nonnegative matrix factorization, joint clustering, visualization, and differential expression testing (Fig. 1A). We carefully compared outputs to ensure that function outputs from the R and Python versions are identical to within the limits of numerical precision. The only exceptions are cases when external packages called by PyLiger, such as UMAP and Leiden, produce slightly different results between R and Python.

As an additional feature, we embedded new functionality for gene ontology (GO) enrichment analysis within PyLiger. This makes it much easier to formulate hypotheses about the functions of key genes that are differentially expressed across cell types or biological conditions. Specifically, PyLiger incorporates GOATOOLS (Klopfenstein *et al*., 2018) for gene ontology enrichment testing and GO-Figure! (Reijnders and Waterhouse, 2021) for visualizing enriched GO terms. For example, given a list of differentially expressed genes, users can easily run a PyLiger function to identify a list of significantly enriched GO terms. They may further visualize the GO terms by semantic similarity scatterplots (Fig. 1A). Functions are fully user-customizable in colormap, labels, etc.

**Fig. 1.**
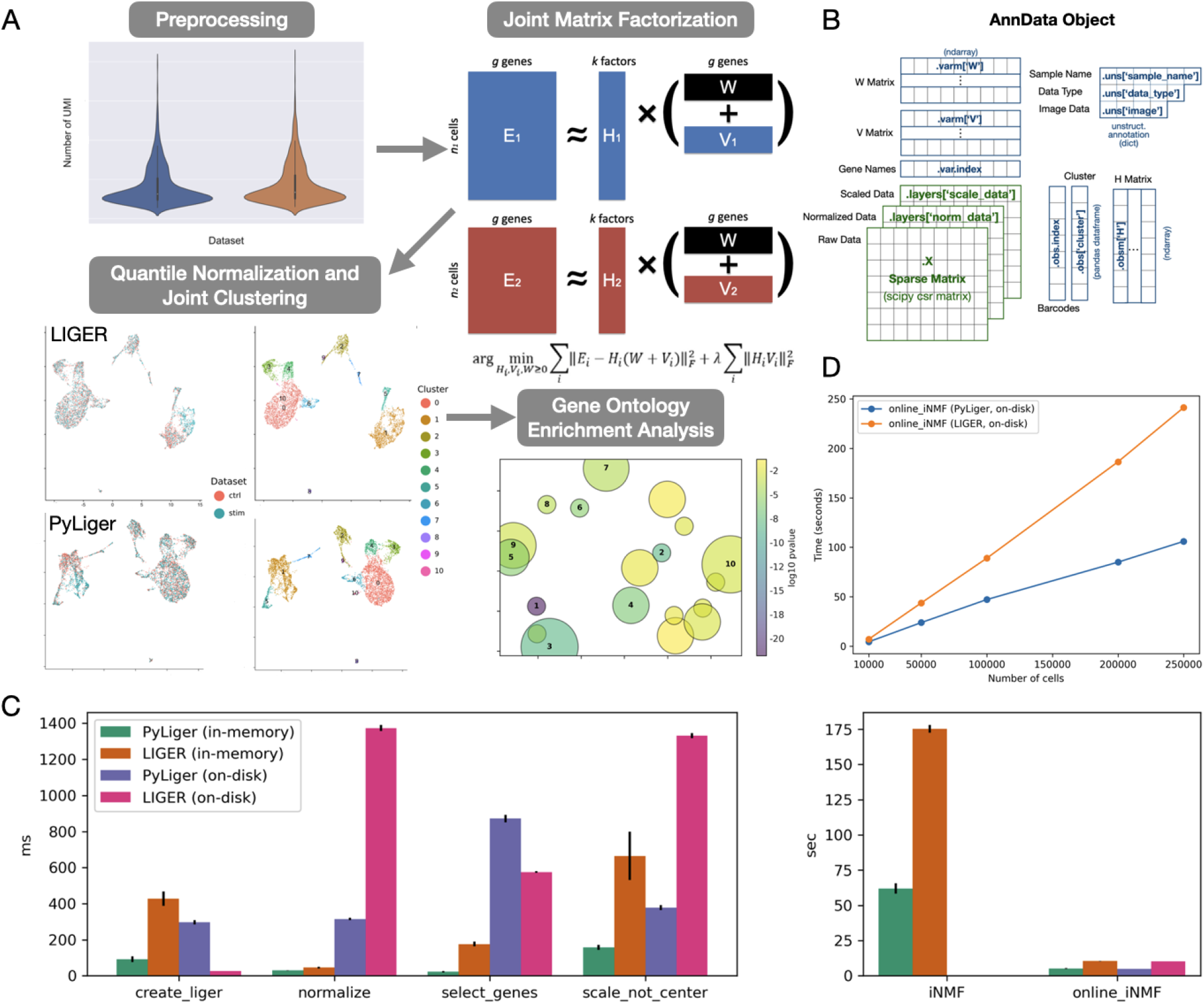
(A) PyLiger functions include preprocessing, joint matrix factorization, joint clustering, visualization, differential expression testing, and GO enrichment analysis. Note that differences in UMAP coordinates are due to differences between R and Python UMAP implementations. (B) Diagram of how Liger class member variables are represented in AnnData format. (C) Comparison of runtimes for PyLiger (in-memory mode), LIGER (in-memory mode), PyLiger (on-disk mode), and LIGER (on-disk mode) functions on a dataset of 6,000 PBMCs. Note that iNMF algorithm only supports in-memory mode, while the online iNMF algorithm can run in on-disk or in-memory mode. (D) Comparison of runtimes for Python and R implementations of online iNMF on a dataset sampled from the adult mouse cortex (n = 10,000, 50,000, 100,000, 200,000 and 255,353 cells datasets)

### 2.2 PyLiger adapts AnnData format to interoperate with existing packages

We designed the structure of the PyLiger class to smoothly interface with the widely used AnnData format. The AnnData package was initially introduced along with Scanpy offering a convenient way to store data matrices and annotations together. We store cell factor loading matrices (*H^i^*), shared metagenes (*W*), and dataset-specific metagenes (*V^i^*) as annotations of the raw matrix (Fig. 1B). The use of AnnData format also facilitates interoperability with existing single-cell analysis tools such as Scanpy and scVelo (Bergen *et al*., 2020). We used the naming rules from Scanpy to name our annotations (UMAP coordinates, for instance) so that each individual AnnData object can be plugged into Scanpy easily.

### 2.3 Python implementation reduces runtimes

To demonstrate the performance of PyLiger package, we tested functions using a dataset of 6,000 PBMCs (Kang *et al*., 2018). We confirmed that the results from PyLiger are identical (to within numerical precision) to those from the LIGER R package. (External packages called by PyLiger, such as UMAP and Leiden, produce slightly different results between R and Python in some cases.) Moreover, PyLiger functions run 1.5 to 5 times faster than their R counterparts (Fig. 1C). In particular, the most time-consuming step–matrix factorization–is approximately 3 times faster in Python than our previous R implementation. This is particularly impressive because many of the R functions are implemented using Rcpp, whereas all of PyLiger is simply implemented in native Python.

### 2.4 PyLiger scales to arbitrarily large single-cell datasets using fixed memory

PyLiger supports HDF5 file format for on-demand loading of datasets stored on disk. We found that in AnnData objects, only raw matrix allows HDF5-based backing, but not other processed matrices stored as layers. Therefore, we store data matrices in a separate HDF5 file while matrix annotations are still stored in AnnData objects. We compared the on-disk mode to the in-memory mode using the same dataset of 6,000 PBMCs. By sacrificing a little processing efficiency (within a second on a dataset of 6,000 cells), the on-disk mode functions can process arbitrarily large datasets using fixed memory (Fig. 1C). Note that the functions *create_liger* and *select_genes* in the on-disk mode of PyLiger are slightly slower than on-disk mode of LIGER due to new feature implementation.

Moreover, we implemented the online iNMF algorithm (Gao *et al*., 2021) in combination with HDF5 file format, providing scalable and efficient data integration as well as significant memory savings. The online iNMF algorithm scales to arbitrarily large numbers of cells but still uses fixed memory and can incorporate new data without recalculating from scratch. To benchmark the performance, we did a comparison of online iNMF between PyLiger and LIGER using five datasets of increasing sizes (ranging from 10,000 to 255,353 cells in total) sampled from the same adult mouse frontal and posterior cortex data. The PyLiger implementation of online iNMF achieves a 2.3 × speedup on average in comparison to its R counterpart (Fig. 1D).

## 3 Conclusion

PyLiger provides an effective way to integrate large-scale single-cell multi-omic datasets. Its Python implementation enables convenient interoperability with other single-cell analysis tools and advanced machine learning and deep learning approaches. Embedded GO enrichment analysis and visualization modules provide a convenient interface for downstream analysis. Furthermore, incorporating online iNMF and HDF5 file format bring PyLiger’s scalability into arbitrarily large numbers of cells.

## Funding

This work was supported by R01HG010883 to JDW.

## Notes

### Competing Interest Statement

The authors have declared no competing interest.

## References

Welch, J. D., Kozareva, V., Ferreira, A., Vanderburg, C., Martin, C., Macosko, E. Z. (2019). Single-cell multi-omic integration compares and contrasts features of brain cell identity. Cell, 177(7), 1873–1887 (2019). https://doi.org/10.1016/j.cell.2019.05.006.

Stuart T, Butler A, Hoffman P, Hafemeister C, Papalexi E, Mauck WM 3rd, Hao Y, Stoeckius M, Smibert P, Satija R. Comprehensive Integration of Single-Cell Data. Cell, 177, 1888–1902 June 13, 2019 a 2019 Elsevier Inc. https://doi.org/10.1016/j.cell.2019.05.031

Korsunsky, I., Millard, N., Fan, J. et al. Fast, sensitive and accurate integration of single-cell data with Harmony. Nat Methods 16, 1289–1296 (2019). https://doi.org/10.1038/s41592-019-0619-0

Wolf, F., Angerer, P. Theis, F. SCANPY: large-scale single-cell gene expression data analysis. Genome Biol 19, 15 (2018). https://doi.org/10.1186/s13059-017-1382-0

Gayoso A, Lopez R, Xing G, Boyeau P, Wu K, Jayasuriya M, Melhman E, Langevin M, Liu Y, Samaran J, Misrachi G, Nazaret A, Clivio O, Xu C, Ashuach T, Lotfollahi M, Svensson V, Beltrame E, Talavera-López C, Pachter L, Theis F. J., Streets A, Jordan M. I., Regier J, Yosef N. scvi-tools: a library for deep probabilistic analysis of single-cell omics data. bioRxiv 2021.04.28.441833; doi: https://doi.org/10.1101/2021.04.28.441833

Klopfenstein, D.V., Zhang, L., Pedersen, B.S. et al. GOATOOLS:APython library for Gene Ontology analyses. Sci Rep 8, 10872 (2018). https://doi.org/10.1038/s41598-018-28948-z

Reijnders MJMF and Waterhouse RM (2021) Summary Visualizations of Gene Ontology Terms With GO-Figure!Front. Bioinform. 1:638255. doi: 10.3389/fbinf.2021.638255

Bergen, V., Lange, M., Peidli, S. et al. Generalizing RNA velocity to transient cell states through dynamical modeling. Nat Biotechnol 38, 1408–1414 (2020). https://doi.org/10.1038/s41587-020-0591-3

Kang, H., Subramaniam, M., Targ, S. et al. Multiplexed droplet single-cell RNA-sequencing using natural genetic variation. Nat Biotechnol 36, 89–94 (2018). https://doi.org/10.1038/nbt.4042

Gao, C., Liu, J., Kriebel, A.R. et al. Iterative single-cell multi-omic integration using online learning. Nat Biotechnol 39, 1000–1007 (2021). https://doi.org/10.1038/s41587-021-00867-x

